# Regression QSAR Models for Predicting HIV-1 Integrase Inhibitors

**DOI:** 10.1101/2021.02.23.432583

**Authors:** Christopher Ha Heng Xuan, Lee Nung Kion, Taufiq Rahman, Hwang Siaw San, Wai Keat Yam, Xavier Chee

**Affiliations:** Faculty of Engineering, Computing and Science, Swinburne University of Technology, Sarawak, Malaysia; Faculty of Cognitive Sciences and Human Development, University Malaysia Sarawak, Sarawak Malaysia; Department of Pharmacology, University of Cambridge, United Kingdom; Centre for Bioinformatics, School of Data Sciences, Perdana University, Selangor Darul Ehsan, Malaysia

## Abstract

The Human Immunodeficiency Virus (HIV) infection is a global pandemic that has claimed 33 million lives to date. One of the most efficacious treatment for naïve or pre-treated HIV patients is with the HIV integrase strand transfer inhibitors (INSTIs). However, given that HIV treatment is life-long, the emergence of HIV-1 strains resistant to INSTIs is an imminent challenge. In this work, we showed two best regression QSAR models that were constructed using a boosted Random Forest algorithm 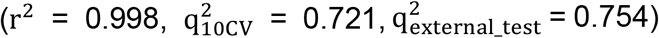 and a boosted K* algorithm 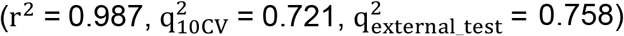 to predict the pIC_50_ values of INSTIs. Subsequently, the regression QSAR models were deployed against the Drugbank database for drug repositioning. The top ranked compounds were further evaluated for their target engagement activity using molecular docking studies and their potential as INSTIs evaluated from our literature search. Our study offers the first example of a large-scale regression QSAR modelling effort for discovering highly active INSTIs to combat HIV infection.

## Introduction

The Human Immunodeficiency Virus (HIV) infection is a global epidemic that have claimed 33 million lives to date. As of 2019, the World Health Organization estimated that 38 million people around the world is living with HIV. HIV infection was one of the leading causes of death in the United States until the development of the Highly Active Anti-retroviral Therapy (HAART).^1^HAART is a combination treatment consisting of several types of drugs, namely reverse transcriptase inhibitors, protease inhibitors, fusion inhibitors, chemokine receptor antagonists and integrase strand transfer inhibitors (INSTIs). The latter is a class of anti-retroviral drug that targets the HIV integrase (IN). IN is the key enzyme involved in incorporating viral DNA into the host CD4 cells through two distinct steps: (a) 3’-processing of the viral DNA to form 3’-OH recessed ends and (b) stabilizing the IN-DNA complex (intasome) for the 3’-OH of the viral DNA to attack the host DNA.^2,3^Currently, there are four FDA-approved INSTIs (raltegravir, elvitegravir, dolutegravir and bictegravir; Figure 1) whilst another agent (cabotegravir; Figure 1) was recently approved by FDA in Jan 2021. Developing INSTIs is an attractive strategy against HIV infection because these drugs have shown high efficacy, exhibit fewer drug-drug interactions and have minimal off-target effects in the human cells.^4,5,6^ However, the long-term use of HIV drugs and the error-prone replication of HIV have given rise to raltegravir- and elvitegravir-resistant HIV strains.^7^ Although second-generation INSTIs like dolutegravir have higher genetic barrier to resistance, the emergence of resistant HIV strains to other INSTIs is not a question of if, but when.^8^ Therefore, there is a pressing need to develop more INSTIs.

**Figure 1.**
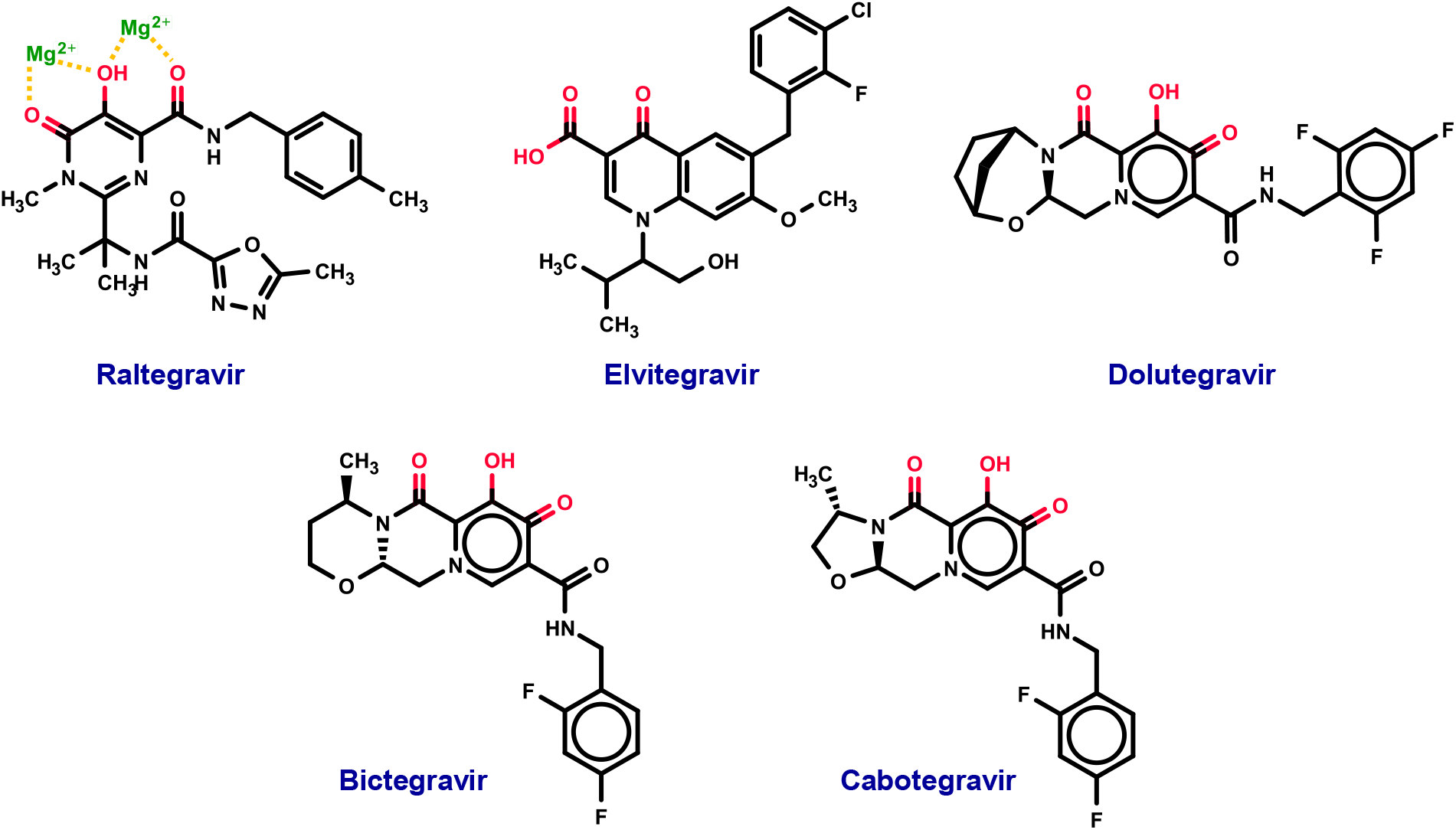
Chemical structures of FDA-approved and experimental INSTIs.

Conventionally, high-throughput screening (HTS) is used to discover lead candidates against a therapeutic target. However, the low hit-rate (0.01-0.1%) and the cost associated with screening millions of compounds renders HTS a very expensive and time-consuming endeavour.^9^ An alternative to conventional HTS can be the experimental screening of compounds that are pre-selected through QSAR-based virtual screening. In developing a QSAR model, molecular descriptors (termed as features) are calculated for a set of known inhibitors. These features are then correlated with the biological activity of these inhibitors, often with the use of machine learning techniques.^9^ Indeed, several regression QSAR models for HIV-1 INSTIs have been reported based on techniques such as molecular docking^10^, comparative field molecular analysis^11^ (COMFA) and comparative molecular similarity indices analysis^12^ (COMSIA). However, these QSAR models are specific to particular classes of HIV-1 INSTIs such as carboxylic acid derivatives (trained on 62 compounds)^13^, curcumine (trained on 29 compounds)^14^, pyridinone (trained on 53 compounds)^15^, *β*-diketo-acids (trained on 37 on compounds)^16^and napthyridine (trained on 50 compounds)^17^. As far as we are concerned, our study is the first large-scale QSAR study (trained on 1417 compounds) that covers a broad chemical structure diversity.

The aim of this study is to establish key compound features that are important for designing next-generation INSTIs. To do this, we constructed two INSTIs regression QSAR models built using the boosted Random Forest^18^ and K* algorithm^19^. The two QSAR models were evaluated using 10-fold cross-validation, external test set and y-randomization test. As part of our drug repositioning effort, these two models were then deployed against the Drugbank compound dataset containing FDA-approved, experimental and investigational drugs to shortlist known drugs that could potentially be repositioned as anti-HIV-1 drugs.

## Results and Discussions

### Preparation of Training Set and Test Set

In this section, we present the results of our regression QSAR models using the workflow showïn in Figure 2. First, we downloaded the compound dataset (CHEMB2366505) that contains approximately 4500 chemical compounds that were experimentally evaluated against IN. After filtering for duplicates, FDA-approved INSTIs and compounds with molecular weight > 680 Da, we were left with 2028 compounds. The bioactivity of these compounds were converted to pIC_50_ – the negative logarithmic value of the concentration required to inhibit 50% strand transfer activity in bioassays. Next, these compounds were then split into training and test sets with a 70:30 split. This resulted in a training set and test set containing 1417 and 611 compounds, respectively. To show that the training and test sets are congruent in terms of chemical similarity, we mapped out the coverage of the chemical space of the compounds in both sets using Principal Component Analysis (PCA). The features used for PCA analysis were the molecular weight, octanol-water partition coefficient (LogP), number of rotatable bonds, total polar surface area (TPSA), number of H-bond donors and number of H-bond acceptors. Using these features, the Principal Components (PCs) 1 and 2 were able to capture 70% of the chemical structure variance. The results are shown in Supplementary Figure S1 and S2.

**Figure 2.**
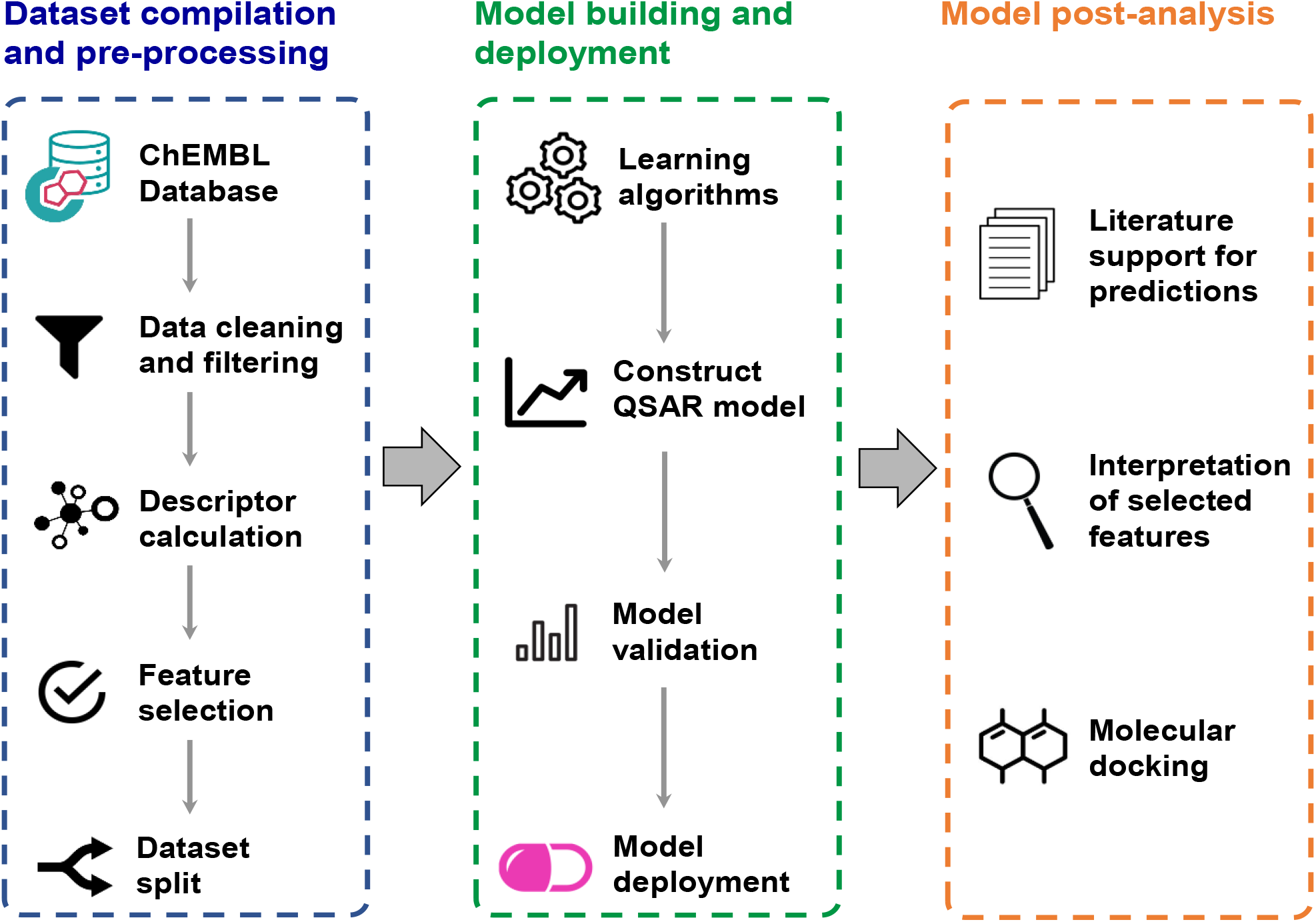
**Graphical scheme of the workflow for constructing regression QSAR models to predict potential INSTIs.**

### Compound featurization and feature selection

Next, the compounds in both training and test sets were featurised using 206 2D-molecular descriptors. To reduce model complexity and prevent data overfit, we used the correlation-based feature selection subset evaluator (CfsSubsetEval) available in Waikato Environment of Knowledge Analysis (WEKA) package^20^. The CfsSubsetEval method evaluates a subset of molecular descriptors by considering their correlation to the pIC_50_ along with the degree of redundancy between the molecular descriptors.^21^ Using this evaluator, 12 molecular descriptors were selected for further regression modelling. The explanation of the molecular descriptors and their correlation to pIC50 are shown in Table 1. To avoid multi-collinearity, we plotted a correlation matrix for the molecular descriptors and showed them in Supplementary Figure S3.

**Table 1.**
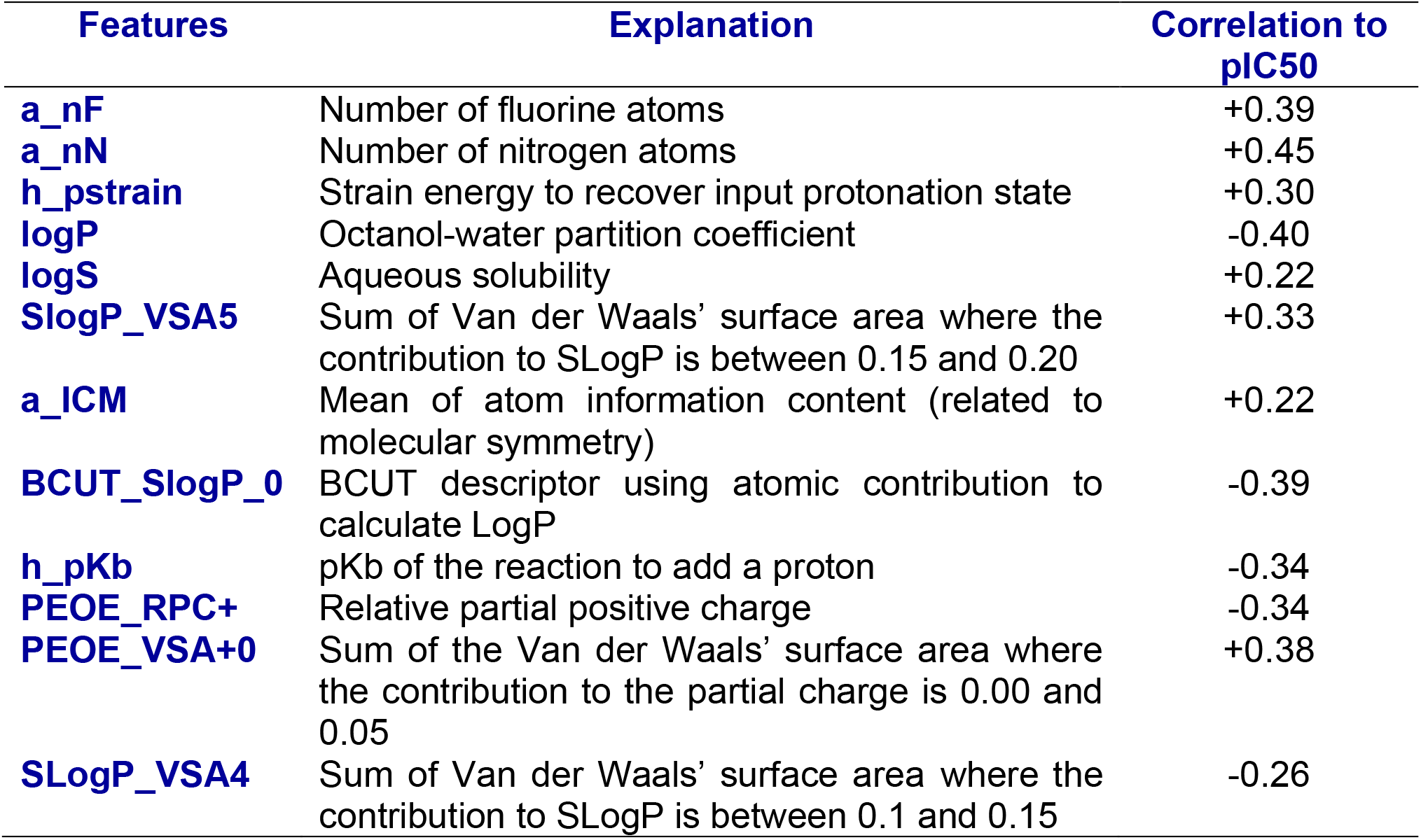
**Explanation and correlation of the molecular descriptors to the experimental pIC**_**50**_ **of compounds.**

| Features | Explanation | Correlation to pIC<sub>50</sub> |
| --- | --- | --- |
| **a_nF** | Number of fluorine atoms | +0.39 |
| **a_nN** | Number of nitrogen atoms | +0.45 |
| **h_pstrain** | Strain energy to recover input protonation state | +0.30 |
| **logP** | Octanol-water partition coefficient | -0.40 |
| **logS** | Aqueous solubility | +0.22 |
| **SlogP_VSA5** | Sum of Van der Waals' surface area where the contribution to SLogP is between 0.15 and 0.20 | +0.33 |
| **a_ICM** | Mean of atom information content (related to molecular symmetry) | +0.22 |
| **BCUT_SlogP_0** | BCUT descriptor using atomic contribution to calculate LogP | -0.39 |
| **h_pKb** | pKb of the reaction to add a proton | -0.34 |
| **PEOE_RPC+** | Relative partial positive charge | -0.34 |
| **PEOE_VSA+0** | Sum of the Van der Waals' surface area where the contribution to the partial charge is 0.00 and 0.05 | +0.38 |
| **SLogP_VSA4** | Sum of Van der Waals' surface area where the contribution to SLogP is between 0.1 and 0.15 | -0.26 |

To understand how these descriptors could inform about drug design for INSTIs, we attempted to correlate some of these descriptors with their biological significance. First, the descriptor PEOE_RPC+ is related to the relative partial positive charge of the inhibitors. This term is negatively correlated with pIC_50_. This is unsurprising given that potential INSTIs are expected to bind and interact with the positively-charged magnesium ions. Besides, the number of fluorine and nitrogen atoms were selected as positive determinants of pIC_50_. Indeed, the very electronegative fluorine and nitrogen atoms are known to alter the physicochemical properties of inhibitors and enhance protein-binding interactions. ^22,23^ Meanwhile, the descriptors LogP, LogS and BCUT_SLogP_0 are related to the solubility of the inhibitors. Next, a negatively correlated h_pKb term indicates that stronger bases are more likely to be INTIs. The descriptors PEOE_VSA+0 is a term related to Van der Waals’ surface area based on electronic properties. The positive correlation of this term indicates that the pIC_50_ of HIV-1 INSTIs is enhanced by their lipophilicity. This is in agreement with experimental SAR studies where increasing lipophilicity of the ring system of HIV-1 INSTIs leads to higher inhibitory activity.^24,25^ Lastly, the descriptors SlogP_VSA5 and SLogP_VSA4 are related to the Van der Waal’s surface area that contributes to a particular range of LogP values while h_pstrain concerns the strain energy to recover the protonation state.

### Regression modelling

We applied four base learners, namely Support Vector Machine (SVM), Random Forest (RF), k-Nearest Neighbour (kNN) and K* algorithms to predict the pIC_50_ of the inhibitors in the training set. Additionally, we also boosted these four base learners using Additive Regression modelling. Additive Regression (AR) modelling is the WEKA’s equivalent to Gradient Boosting for enhancing the performance of regression modelling algorithms. To prevent over-fitting, we evaluated all algorithms using 10-fold cross validation (CV) too. From our study, we obtained a highly predictive AR model with RF as the base learner. Besides, another comparable regression model was constructed using AR model with the K* algorithm. Of note, while k-NN as a base learner (with or without AR) performed well when evaluated on the training set, its performance dropped when tested using the 10-fold CV. Hence, k-NN was discarded from further study. The performance of these base learners (with or without boosting) is summarized in Supplementary Table ST1.

Subsequently, we search for the hyperparameter for both the AR-RF and AR-K* algorithms to improve their performance in regression modelling. The tuned AR-RF and AR-K* were then re-evaluated using 10-CV and the previously constructed external test set. The performance for the tuned AR-RF and AR-K* is summarized in Supplementary Table ST2. The scatter plots for the experimentally obtained pIC50 and predicted pIC50 as well as the plots of the pIC_50_ residuals for both AR-RF and AR-K* are shown in Figure 3a-c and Figure 4a-c, respectively. To study (a) the occurrence of chance correlation and (b) responsiveness of pIC_50_ to the selected features, we conducted y-randomization test. In this test, the pIC50 of the compounds in the training sets were randomly shuffled and new QSAR models were created. In each of our 20 randomization runs for both AR-RF and AR-K*, we observed that the r^2^ of regression models built using y-randomized training sets were above 0.9. This indicated that our machine learning algorithms were prone to overfitting the data and this phenomenon was similarly observed by Darnag *et al*.^26^ However, none of the random regression QSAR models had any predictability power when evaluated using the external test set (Figure 3d and 4d).

**Figure 3.**
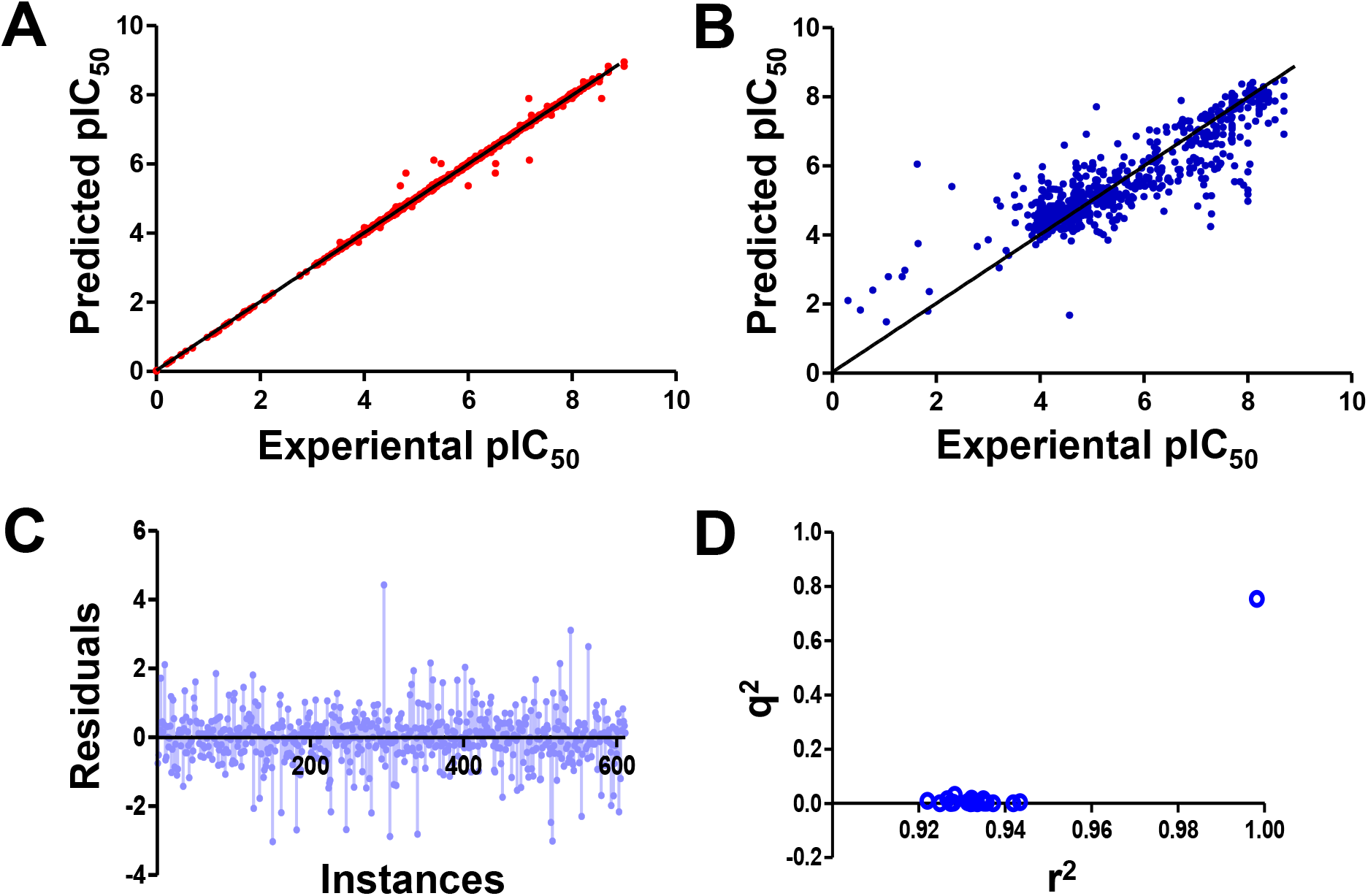
Performance of regression QSAR model constructed using AR-RF algorithm. **(A)** Graph plot of predicted pIC_50_ against experimental pIC_50_ of training set (1417 compounds). **(B)** Graph plot of predicted pIC_50_ against experimental pIC_50_ of test set (611 compounds). **(C)** Residual plot of the predicted pIC_50_ and the experimental pIC_50_ values of the compounds in the test set. **(D)** Y-randomization plot for the regression model evaluated on test set (20 randomized models)

**Figure 4.**
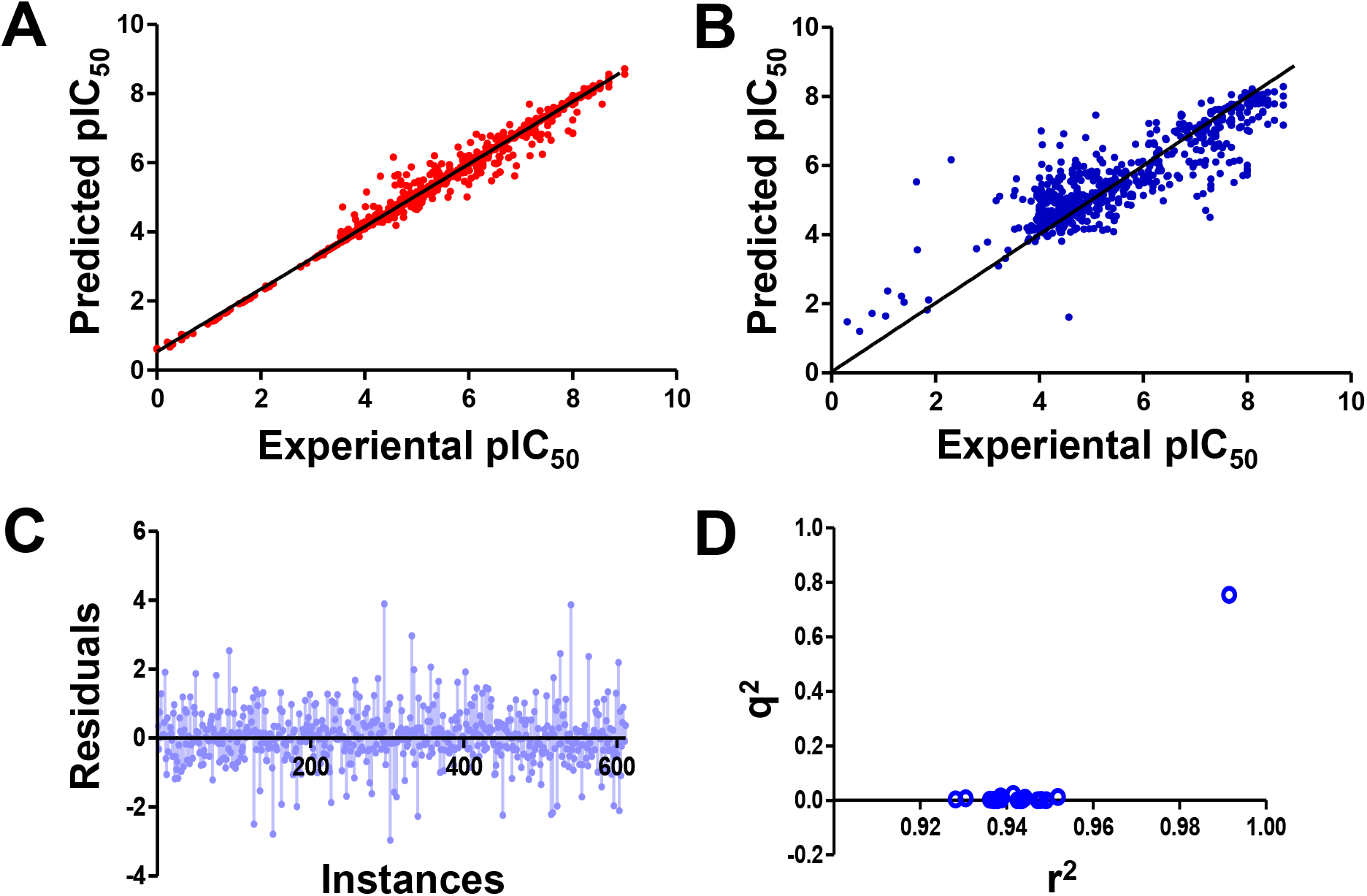
Performance of regression QSAR model constructed using AR-K* algorithm. **(A)** Graph plot of predicted pIC_50_ against experimental pIC_50_ of training set (1417 compounds). **(B)** Graph plot of predicted pIC_50_ against experimental pIC_50_ of test set (611 compounds). **(C)** Residual plot of the predicted pIC_50_ and the experimental pIC_50_ values of the compounds in the test set. **(D)** Y-randomization plot for the regression model evaluated on test set (20 randomized models)

In general, statistical values of r^2^ > 0.6 and q^2^ > 0.5 between the predicted and experimental value indicated good predictability for the QSAR models. Judging from this, we have decided to deploy both AR-RF and AR-K* models to predict potential HIV-1 INSTIs.

### Model deployment

We deployed both AR-RF and AR-K* on the Drugbank database, which contains FDA-approved, experimental and investigational drugs. We selected the Drugbank database for screening because we were intrigued with the idea of repositioning known or experimental drugs to target HIV IN. Drug repositioning could speed up drug development as repositioned drugs have gone through extensive pharmacokinetic studies and are less likely to fail in clinical trials due to toxicity effects. From our screening, it is interesting to note that known INSTIs such as the experimental drug GSK364735, Dolutegravir, Bictegravir and Cabotegravir were ranked among the top five chemicals predicted to be HIV-1 INSTIs by AR-RF. We showed the chemicals predicted to be HIV-1 INSTIs in Figure 5 and Table 2. After screening, we pooled the predicted pIC50 of the chemicals by AR-RF and AR-K* for consensus ranking. For chemicals to be considered as potential HIV-1 INSTIs, their predicted pIC50 must fall above the top 75% quartile in both regression model. The full list of chemicals predicted to be HIV-1 INSTIs based on our criterion are attached in Supplementary Table ST3.

**Table 2.** The predicted pIC50 of the top-scoring compounds as ranked by the AR-RF regression model. Experimental and FDA-approved HIV-1 INSTIs are highlighted with red asterisks.

| Rank | Common drug name | Drugbank ID | Predicted pIC50 from AR-RF | Predicted pIC50 from AR-K* |
| --- | --- | --- | --- | --- |
| 1 | GSK-364 735* | DB13119 | 8.101 | 8.116 |
| 2 | Dolutegravir* | DB08930 | 7.896 | 7.693 |
| 3 | Itacitinib | DB12154 | 7.603 | 7.059 |
| 4 | Bictegravir* | DB11799 | 7.553 | 7.682 |
| 5 | Cabotegravir* | DB11751 | 7.482 | 7.700 |
| 6 | Rosuvastatin | DB01098 | 7.319 | 7.285 |
| 7 | Verubecestat | DB12285 | 7.296 | 7.382 |
| 8 | - | DB07092 | 7.201 | 7.579 |
| 9 | Epacadostat | DB11717 | 7.184 | 7.779 |
| 10 | - | DB03202 | 7.100 | 7.525 |

**Figure 5.**
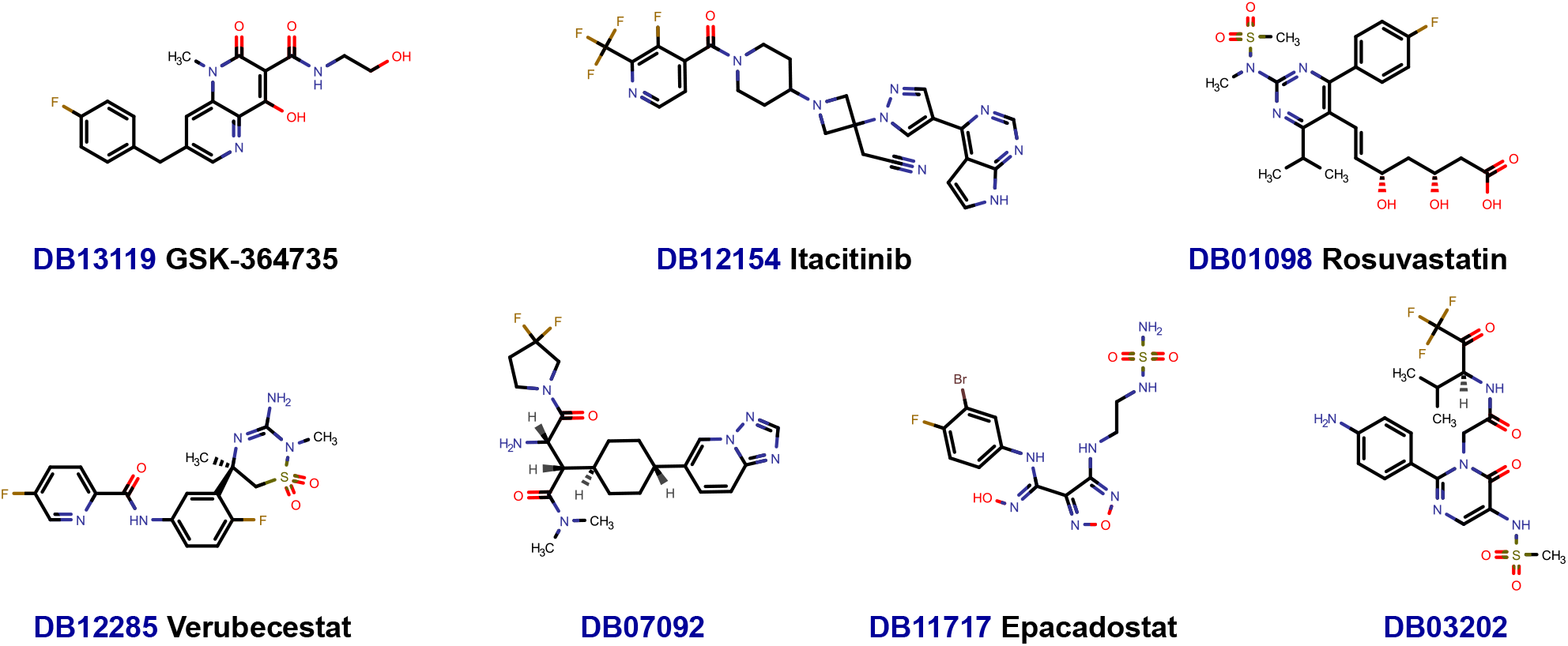
Chemical structures of top compounds predicted to be INSTIs.

To further solidify our confidence in the predicted compounds, we looked into the literature if any of the chemicals in the list were reported to have anti-HIV-1 activity.

Itacitinib is an experimental Janus kinase-1 (JAK1) inhibitor with immunomodulatory activity for treatment against acute graft-versus-host disease.^27^ Although itacitinib was not reported to have anti-HIV-1 activity, it is interesting to note that two structurally similar JAK1/2 inhibitors – Ruxolitinb and Tofacitinib – are reported to demonstrate sub-micromolar inhibition of infection of HIV-1, HIV-2 and a chimeric simian-human immunodeficiency virus carrying reverse transcriptase (RT-SHIV). One study showed that JAK 1/2 inhibitors could down-regulate HIV-induced inflammation that favours viral replication and disease progression.^28^

Rosuvastatin is an FDA-approved HMG-CoA reductase inhibitor (also known as statins) that lowers cholestrol levels.^29^ Aside from their lipid-lowering activity, *in vitro* studies have shown that statins could also exhibit anti-HIV activity. The anti-HIV activity of statins are suggested to stem from down-modulation of lipid rafts necessary for HIV infection in host cell^30,31^, down-regulating Rho activity^32^ or by blocking the integrin intercellular adhesion molecule 1 (ICAM) on the host cell surface to prevent viral entry^33^. Given the range of pleitropic effects that statins exhibit, it is also plausible that statins could exhibit anti-HIV effects through inhibition of HIV IN. Additionally, the chemical structure of rosuvastatin resembles the metal-chelating diketo acid moiety of raltegravir.

Verubecestat is an experimental beta-secretase (BACE1) inhibitor for treatment of Alzheimer ‘s disease.^34^ Meanwhile, epacadostat is an inhibitor of indoleamine-2,3-dioxygenase (IDO1) with anti-neoplastic activity. Although both drugs were not reported directly to have HIV-1 activity, the N-phenylpicolinamide moiety of verubecestat and the 1,2,5-oxadiazole moiety of epacadostat were reported to carry inhibitory activity against HIV-1 IN.^35,36,37^

### Molecular docking

Following on, we selected the top three compounds – itacitinib, rosuvastatin and verubecestat – for molecular docking studies to explore the binding interactions of these compounds with HIV-1 IN. As there is no full-length HIV-1 IN crystallized to-date, we chose to conduct our molecular docking studies using the Prototype Foamy Virus (PFV) IN as a proxy (PDB ID: 3OYA). We chose PFV IN as the model on the basis that the regions near the active sites of the PFV and HIV IN are highly conserved due to similar residues involved in substrate binding and catalysis.^38^ These residues involve in the PFV In catalytic triads are Glu211, Asp128 and Asp185. The corresponding residues in HIV-1 in are Glu152, Asp64 and Asp116. Indeed, the high degree of similarity between the PFV and HIV-1 IN has prompted another study to conclude that the PFV IN was a more accurate model for virtual screening studies for INTIs compared to HIV-1 IN homology models.^39^ Additionally, the co-crystallized viral DNA and raltegravir in the PFV intasome would allow further understanding of protein-inhibitor interaction.

First, we conducted cognate docking study by re-docking raltegravir into the PFV IN. The purpose of cognate docking is to validate pose prediction quality of the GOLD molecular docking software. In the cognate docking study, GOLD was largely able to reproduce the co-crystallized pose of raltegravir (pose root mean square deviation = 0.285Å) with the exception of the 1,3,4-oxadiazole ring. The result of the cognate docking study is in Supplementary Figure S4. Next, we docked itacitinib, rosuvastatin and verubcestat into the PFV IN. The predicted binding poses of these compounds are illustrated in Figure 6. In all three cases, the inhibitors were predicted to coordinate to the catalytic magnesium ions in the active site of the IN enzyme. The coordinating distances of itacitinib, rosuvastatin and verubcestat to the magnesium ions were between 1.8-2.3Å as measured using PyMOL. This is an important observation as several studies have alluded to the metal-dependent inhibition of HIV-1 IN.^40,41,42^ Additionally, all three inhibitors also interacted with the PFV IN catalytic triad in a similar manner to raltegravir. For example, all three inhibitors exhibited magnesium ion-mediated bonding with Asp128. With the exception of itacitinib, both rosuvastatin and verubecestat were involved in another metal-mediated bonding with Asp185. Other noteworthy interactions include the interaction with Tyr212 (Tyr143 in HIV-1 IN) by itacitinib and rosuvastatin and the interaction with Pro214 (Pro145 in HIV-1 IN) by all three inhibitors. These two interactions are important for the stabilization of raltegravir in the active site.^43^ The protein-inhibitor interaction diagrams are in Supplementary Figure S5-S7

**Figure 6.**
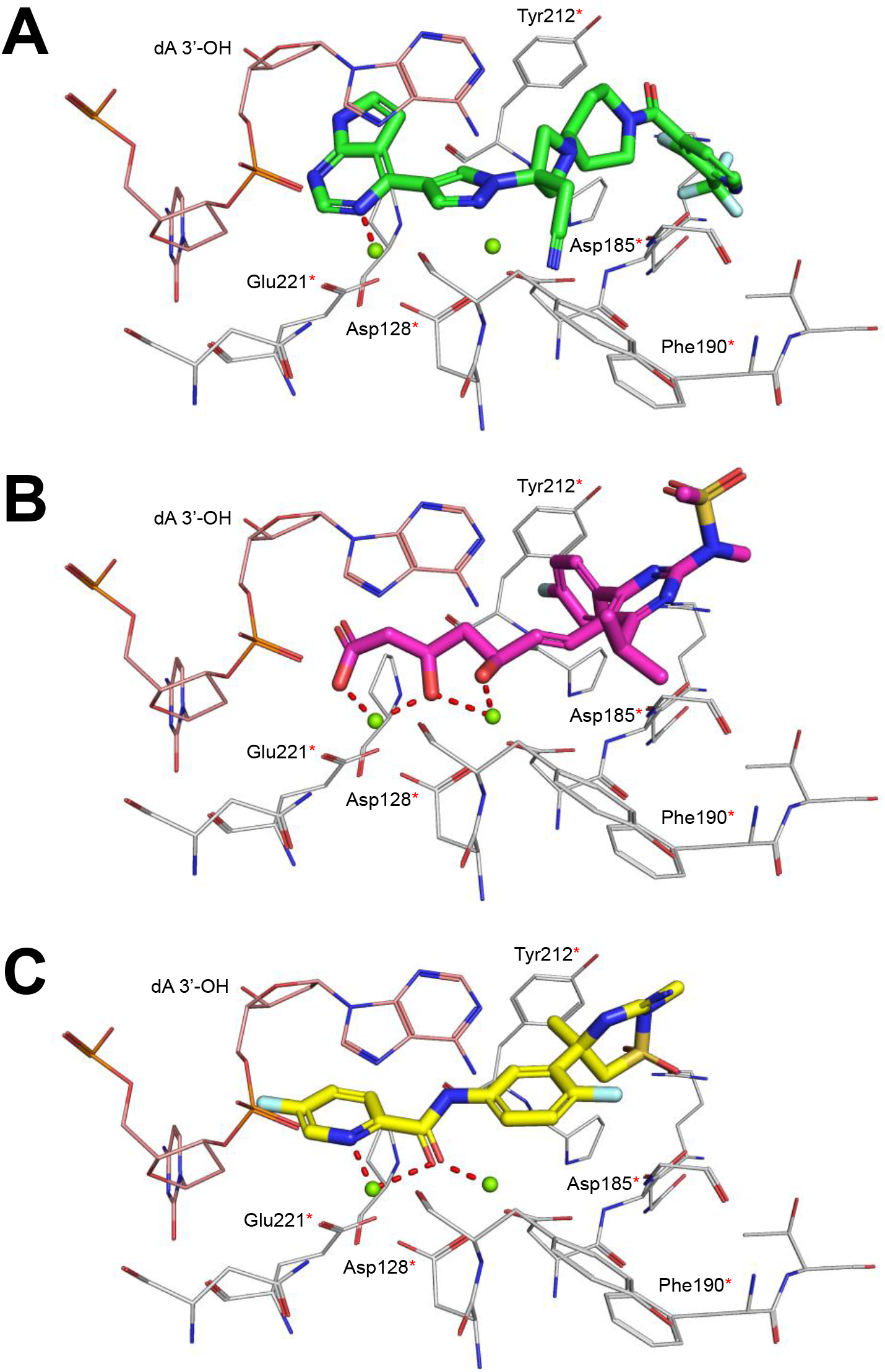
Binding interactions of the PFV IN active site with the top ranked compounds. The docked pose of **(A)** itacitinib, **(B)** rosuvastatin and **(C)** verubecetat are shown interacting with the catalytic magnesium ions. Conserved residues found in both HFV and HIV-1 IN active sites are highlighted with red asterisks. The residues of the active site shown as grey lines. The viral DNA shown as light orange lines. Magnesium ions shown in green.

## Conclusion

HIV-1 IN is an attractive therapeutic target because of their high efficacy and lack of structural orthologs in humans. HIV-1 strains that are resistant towards current INSTIs necessitates putting more potential INSTIs in the drug discovery pipeline. To this end, we constructed two regression QSAR models using the RF and K* algorithms that were boosted by AD modelling. Next, we subsequently deployed our regression models for drug repositioning using the Drugbank compound dataset. We selected three top scoring compounds – itacitinib, rosuvastatin and verubecestat – for molecular docking to study the protein-inhibitor interaction. We further suggest that these three compounds have metal-chelating functional groups that could inhibit HIV-1 IN strand transfer activity. As far as we are concerned, this is the first large-scale QSAR study capable of predicting the pIC_50_ of potential INSTIs. Although our study was geared on discovering potential INSTIs through drug repositioning, we believe this approach can be generalized and applied to other biological targets. This would reduce the time taken in the drug discovery process – from molecular conception to having a marketable drug.

## Experimental Methods

### Preparation of Dataset

The HIV-1 integrase inhibitor dataset were downloaded from CHEMBL2366505. Compounds with half maximal inhibitory concentration (IC_50_) were chosen for this study. The IC50 was converted to pIC50 using a negative logarithmic transformation. After removing duplicates and known FDA-approved HIV-1 integrase inhibitors from the dataset, compounds with molecular weight more than 680 Da were removed from the dataset. After processing, a total of 2028 compounds remained. These compounds were neutralized and energy minimized using the Molecular Operating Environment (MOE) 2015.10 package using the default settings. The compounds were then featurized by the 2-dimensional (2D) molecular descriptors by MOE. Next, these compounds were then split by a 70:30 ratio into a training set and a test set for QSAR modelling using machine learning. To ensure that the structures in the training and test sets are similar, the chemical space of the compounds in both datasets were visualized by means of Principal Component Analysis using the Platform for Unified Molecular Analysis (PUMA)^44^.

### QSAR Modelling and Model Deployment

WEKA is a suite containing different machine learning algorithms.^20^ First, important features were selected using the WEKA attribute selector “CfsSubsetEval” with “BestFirst” as the selection method. The selected features were then checked for multi-collinearity using a correlation matrix. Subsequently, the datasets containing the selected features were used as input for the machine algorithms Sequential Minimal Optimization Regression (SMOreg), Random Forest (RF), k-Nearest Neighbour (kNN) and the K* algorithm (with or without boosting from Additive Regression). Boosted RF and K* algorithms were selected as the best performing and their hyperparameters were fine-tuned. The performance of these two algorithms were evaluated using a 10-fold CV and an external test set. The metrics used were q^2^ and RMSE. Y-randomization tests were conducted to evaluate the risk of chance correlation and to quantify the sensitivity of the compound pIC50 values to the selected features. The tests were conducted by randomizing the pIC50 values without modifying the values of the molecular descriptors. Finally, the models were deployed against the Drugbank dataset version 5.1.8.^45^

### Molecular Docking

The crystal structure of the Prototype Foamy Virus (PFV) integrase complexed with viral DNA (PDB ID: 3OYA) was obtained from the RCSB Protein Data Bank. Hydrogens were added to the intasome complex and energy minimized using the AMBER10:EHT forcefield from the MOE package. The viral DNA was kept inside the intasome as inhibitors of integrase are expected to form *π-π* interaction with the base of the viral DNA. The chemical structure of raltegravir, itacitinib, rosuvastatin and verubecestat were drawn using MarvinSketch version 21.1 and energy-minimized using the MOE package. The compounds were docked to the PFV integrase by using the Genetic Optimization for Ligand Docking (GOLD) 5.3.0 package developed by the Cambridge Crystallographic Data Centre (CCDC). The binding site was defined as 10Å radius sphere centred around the raltegravir. All four inhibitors were docked with 50 Genetic Algorithm runs while keeping all other settings to their default. The ChemPLP was used as the scoring function. The protein-ligand interactions of the top-scoring sdocked poses for these compounds were visualized using PyMol 2.0 and LigPlot+ version 2.2.

## Acknowledgements

This work is kindly supported by the Swinburne Strategy Grant (SSRG2-5624) by Swinburne University of Technology (Sarawak Campus) awarded to X.C.

## Author Contributions

X.C. and L.N.K. conceived the study, guided the experimental design and drafted the manuscript. C.H. conducted the QSAR modelling, data collection. T.R. conducted molecular docking studies. H.S.S. and Y.W.K. provided input on data analysis. All authors helped by providing suggestions and ideas for improving the study, and reviewed and approved the submitted the manuscript.

## Additional Information

The author(s) declare no competing interests.

