## Supplementary Figures S1-S7 for "Regression QSAR Models for Predicting HIV-1 Integrase Inhibitors"

### Supplementary Figure S1

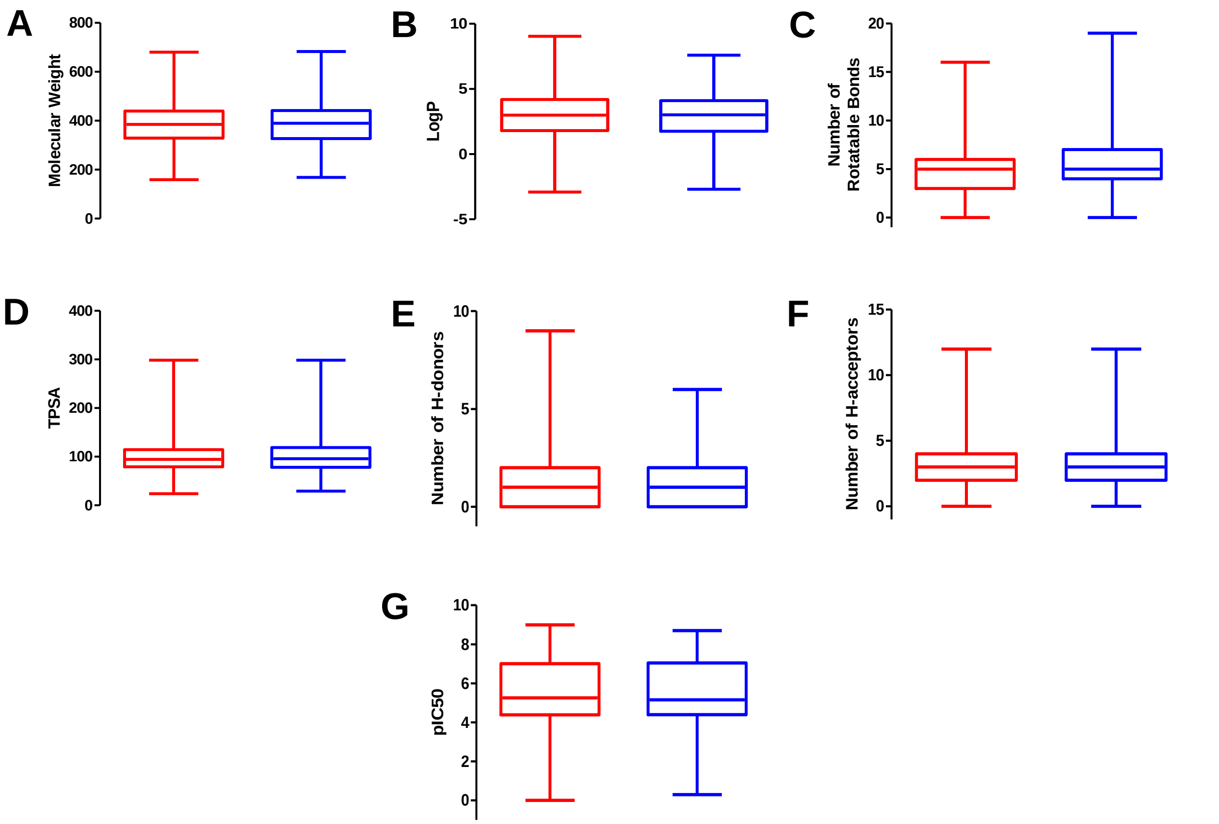


**Figure S1.** Box plots showing distributions of the six features and the $pIC_{50}$ of the compounds in the training set and test set. Training set shown in red; test set shown in blue.

Supplementary Figure S2

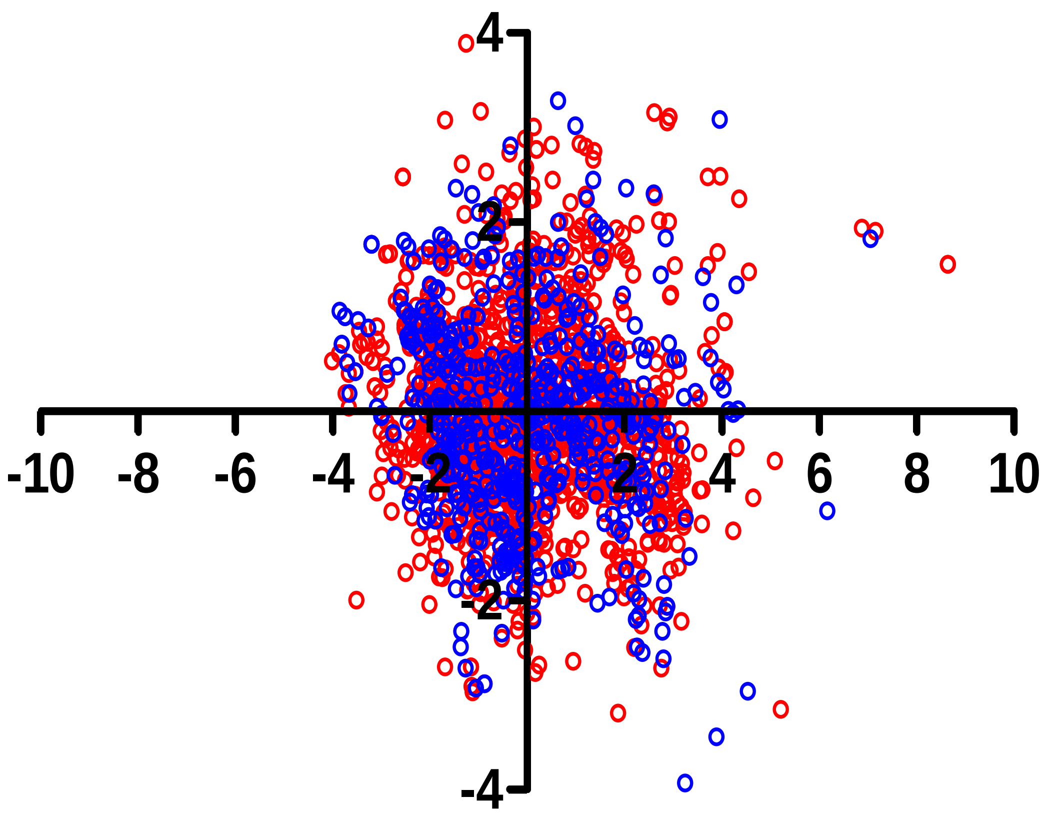


**Figure S2. PCA biplot mapping the chemical space of compounds in the training set and test sets.**

#### Supplementary Figure S3

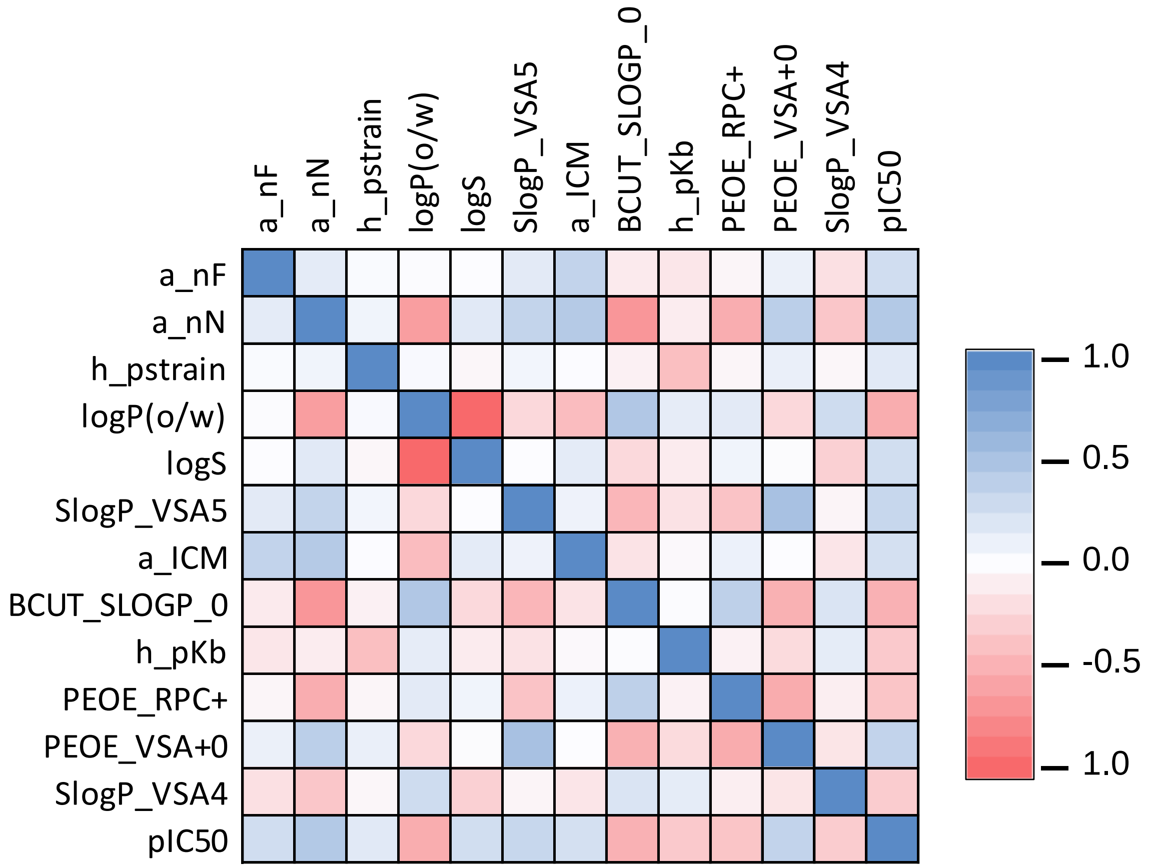


**Figure S3. Correlation matrix showing the degree of collinearity between the selected features and with pIC$_{50}$.**

### Supplementary Figure S4

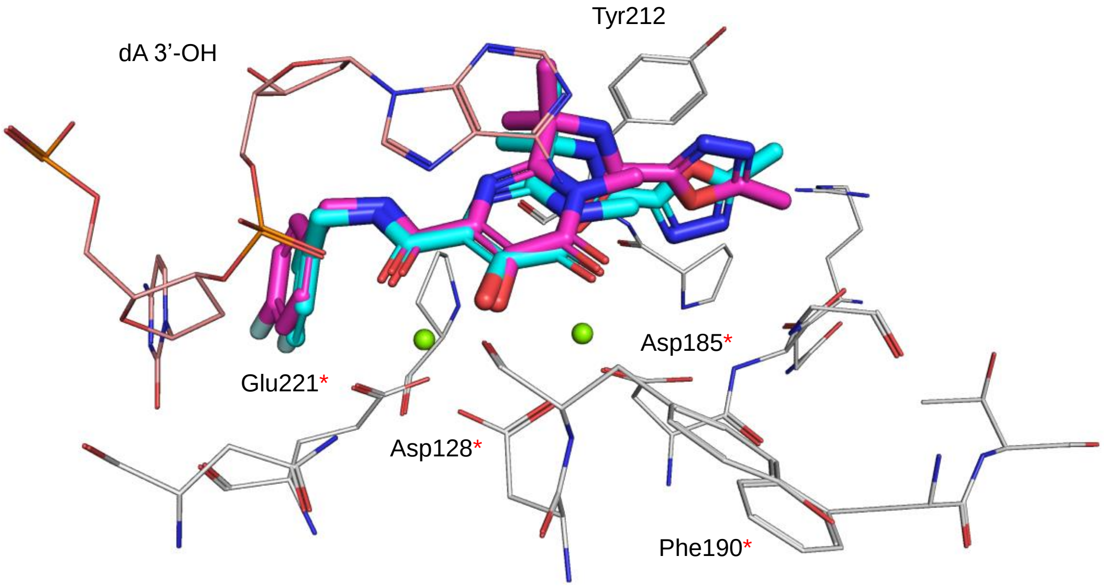


**Figure S4. Cognate docking of raltegravir to PFV IN.** Co-crystallized pose of raltegavir shown in magenta; docked pose predicted by GOLD shown in cyan.

### Supplementary Figure S5

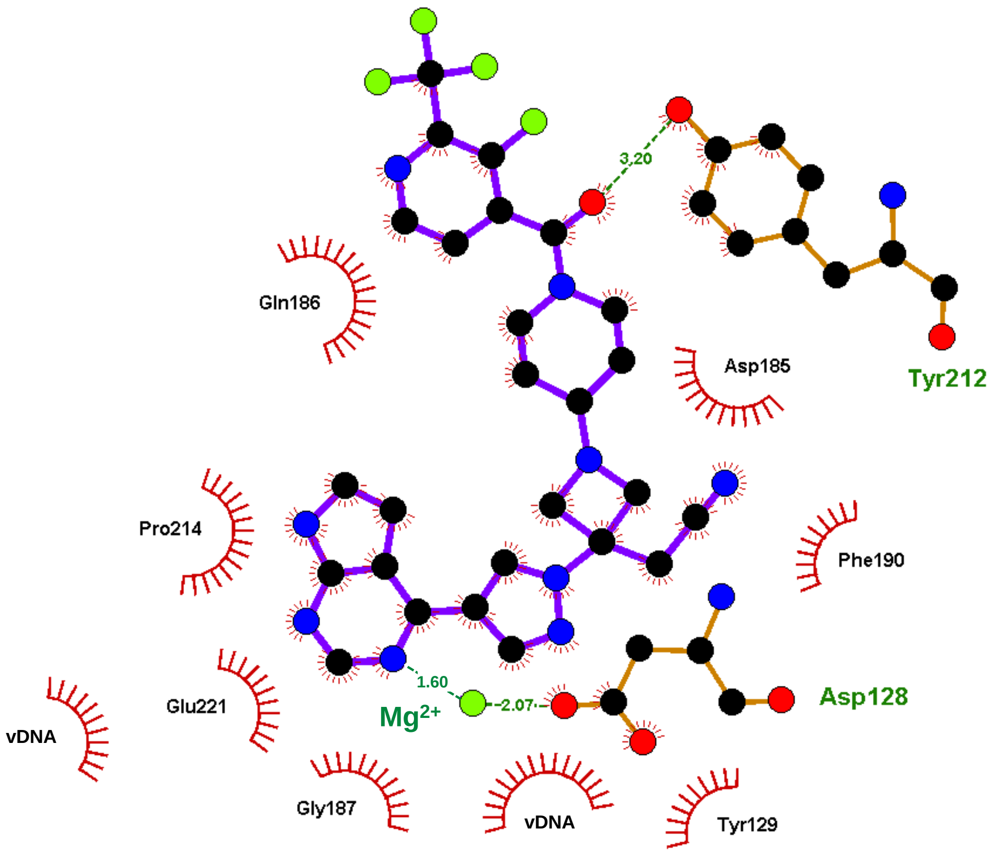


**Figure S5. Interaction of itacitinib with protein residues in the PFV IN active site.** Interaction diagram generated using LigPlot+ developed by the European Bioinformatics Institute. Itacitnib and important interacting residues (Asp128 and Tyr212) shown as balls-and-sticks. Hydrogen bond interactions shown in doted lines with annotated distance in Angstrom. Hydrophobic interactions shown as red fringes. Magnesium ion shown as green sphere.

### Supplementary Figure S6

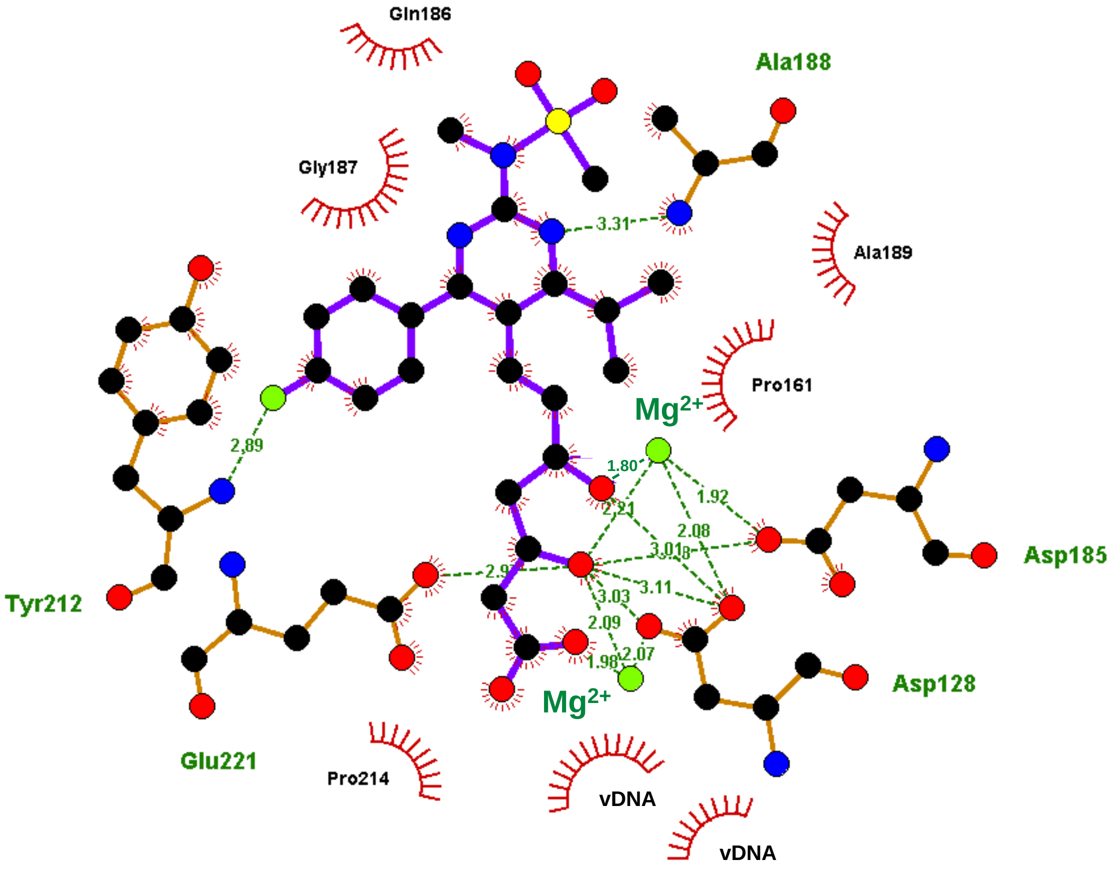


**Figure S6. Interaction of rosuvastatin with protein residues in the PFV IN active site.** Interaction diagram generated using LigPlot+ developed by the European Bioinformatics Institute. Rosuvastatin and important interacting residues (Tyr212, Glu221, Asp128 and Asp185) shown as balls-and-sticks. Hydrogen bond interactions shown in doted lines with annotated distance in Angstrom. Hydrophobic interactions shown as red fringes. Magnesium ions shown as green spheres.

### Supplementary Figure S7

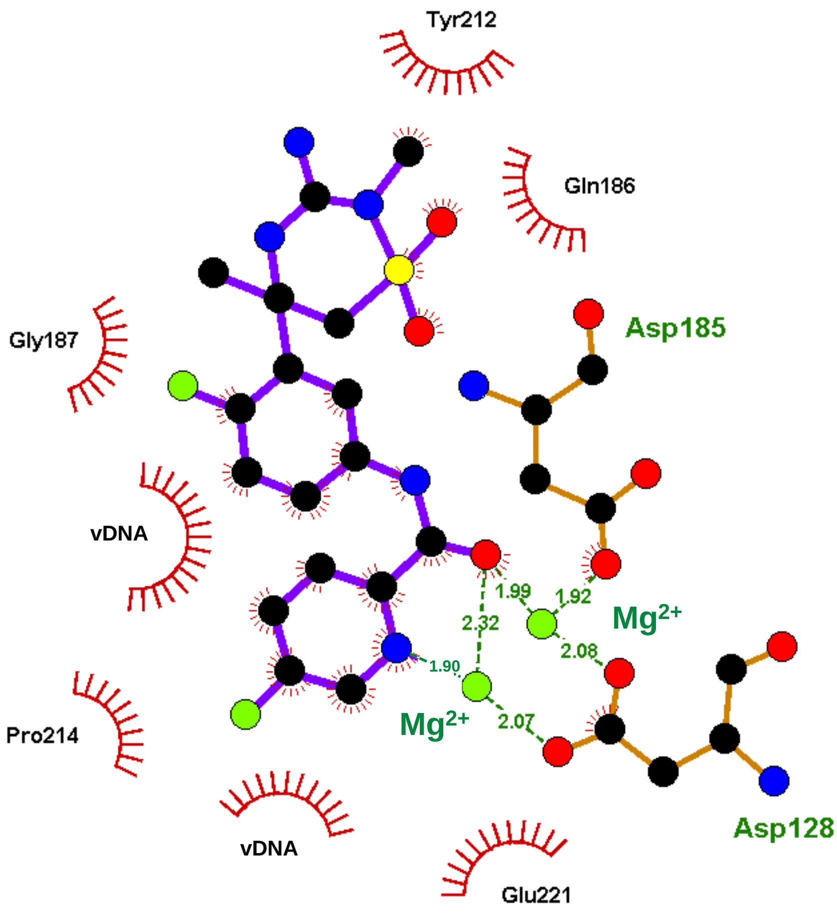


**Figure S7. Interaction of verubecestat with protein residues in the PFV IN active site.** Interaction diagram generated using LigPlot+ developed by the European Bioinformatics Institute. Rosuvastatin and important interacting residues (Asp128 and Asp185) shown as balls-and-sticks. Hydrogen bond interactions shown in doted lines with annotated distance in Angstrom. Hydrophobic interactions shown as red fringes. Magnesium ions shown as green spheres.
