## Supplementary Tables ST1-ST3 for "Regression QSAR Models for Predicting HIV-1 Integrase Inhibitors"

### Supplementary Table ST1

| | AR-RF | | | AR-K* | | |
|---|---|---|---|---|---|---|
| | Training Set | 10-fold CV | Test Set | Training Set | 10-fold CV | Test Set |
| No. of compounds | 1417 | 1417 | 611 | 1417 | 1417 | 611 |
| r / q | 0.999 | 0.849 | 0.868 | 0.993 | 0.849 | 0.870 |
| $r^2$ / $q^2$ | 0.998 | 0.721 | 0.754 | 0.987 | 0.721 | 0.758 |
| $r^2$_adjusted | 0.998 | 0.718 | 0.748 | 0.986 | 0.718 | 0.752 |
| MAE | 0.025 | 0.612 | 0.540 | 0.156 | 0.598 | 0.538 |
| RMSE | 0.069 | 0.863 | 0.775 | 0.227 | 0.865 | 0.771 |

**Supplementary Table 1. Performance of the regression QSAR models constructed using RF and K* algorithms boosted by AD modelling.**

### Supplementary Table ST2

| | Without regression boosting | | | | With regression boosting | | | |
|---|---|---|---|---|---|---|---|---|
| | SVM | RF | kNN | K* | SVM | RF | kNN | K* |
| **Training Set** | | | | | | | | |
| r | 0.624 | 0.983 | 0.999* | 0.995 | 0.624 | 0.999* | 0.999* | 0.997 |
| $r^2$ | 0.389 | 0.967 | 0.998* | 0.990 | 0.389 | 0.998* | 0.998* | 0.994 |
| MAE | 0.974 | 0.237 | 0.007* | 0.049 | 0.974 | 0.009 | 0.007* | 0.025 |
| RMSE | 1.278 | 0.332 | 0.064* | 0.161 | 1.278 | 0.065 | 0.064* | 0.115 |
| **10-fold Cross Validation** | | | | | | | | |
| q | 0.613 | 0.837* | 0.782 | 0.824 | 0.613 | 0.847* | 0.782 | 0.821 |
| $q^2$ | 0.376 | 0.700 | 0.611 | 0.679 | 0.376 | 0.717 | 0.611 | 0.674 |
| MAE | 0.991 | 0.649 | 0.700 | 0.622* | 0.992 | 0.616* | 0.700 | 0.629 |
| RMSE | 1.294 | 0.899* | 1.007 | 0.953 | 1.294 | 0.869* | 1.077 | 0.962 |

**Supplementary Table 2. Performance of different base learners (with or without boosting by additive regression modelling. Top performers of each evaluation metric are highlighted with red asterisks.**

### Supplementary Table ST3

| No | Drugbank ID | Common drug name | AR-RF pIC50 | AR-K* pIC50 | ChemPLP Docking Score |
|---|---|---|---|---|---|
| 1 | DB13119 | GSK-364735* | 8.101 | 8.116 | 73.19 |
| 2 | DB08930 | Dolutegravir* | 7.896 | 7.693 | 60.4 |
| 3 | DB12154 | Itacitinib | 7.603 | 7.059 | 70.87 |
| 4 | DB11799 | Bictegravir* | 7.553 | 7.682 | 66.1 |
| 5 | DB11751 | Cabotegravir* | 7.482 | 7.7 | 63.76 |
| 6 | DB01098 | Rosuvastatin | 7.319 | 7.285 | 72.52 |
| 7 | DB12285 | Verubecestat | 7.296 | 7.382 | 57.35 |
| 8 | DB07092 | - | 7.201 | 7.579 | 73.88 |
| 9 | DB11717 | Epacadostat | 7.184 | 7.779 | 82.69 |
| 10 | DB03202 | - | 7.1 | 7.525 | 68.04 |
| 11 | DB11902 | Gisadenafil | 7.059 | 7.686 | 78.43 |
| 12 | DB11904 | Flumatinib | 7.057 | 6.628 | 74.12 |
| 13 | DB09335 | Alatrofloxacin | 7.049 | 6.852 | 67.72 |
| 14 | DB05294 | Vandetanib | 7.047 | 7.271 | 56.5 |
| 15 | DB07744 | - | 7.031 | 8.014 | 74.57 |
| 16 | DB12720 | Nivocasan | 7.016 | 7.174 | 60.19 |
| 17 | DB12960 | INCB-9471 | 6.991 | 6.225 | 56.15 |
| 18 | DB07781 | - | 6.956 | 7.309 | 57.4 |
| 19 | DB11747 | Barasertib | 6.946 | 6.993 | 101.57 |
| 20 | DB12501 | ABT-384 | 6.92 | 7.261 | 64.08 |
| 21 | DB12130 | Lorlatinib | 6.917 | 7.455 | 68.51 |
| 22 | DB12241 | TAK-733 | 6.912 | 7.307 | 70.41 |
| 23 | DB07665 | - | 6.905 | 7.235 | 82.95 |
| 24 | DB12904 | ZSTK-474 | 6.894 | 7.389 | 65.7 |
| 25 | DB08403 | - | 6.825 | 8.032 | 73.19 |
| 26 | DB06652 | Vicriviroc | 6.823 | 5.839 | 58.76 |
| 27 | DB12479 | Zabofloxacin | 6.819 | 6.971 | 51.58 |
| 28 | DB04591 | - | 6.809 | 6.794 | 71.12 |

**Supplementary Table 3. Compounds from the Drugbank dataset predicted as potential HIV-1 INSTIs.** Experimental and FDA-approved HIV-1 INSTIs are highlighted with red asterisks.
